# Stabilizing selection generates selection against introgressed DNA

**DOI:** 10.1101/2024.08.20.608860

**Authors:** Carl Veller, Yuval Simons

## Abstract

DNA introgressed from one population into another is often deleterious to the recipient population if the two populations have diverged genetically from one another. Previous explanations of this phenomenon have posited negative interactions between donor-population alleles and the recipient population’s genome or environment, or higher genetic load in the donor population. Here, we show that stabilizing selection on quantitative traits—even around the same optimal trait values in the two populations and when the populations are demographically identical—generates selection against the minor-parent ancestry in a population formed via unequal admixture of the two populations. We calculate the rate at which minor-parent ancestry is purged under this mechanism, both in the early generations after admixture and in the long term, and we verify these calculations with whole-genome simulations. Because of its ubiquity, stabilizing selection offers a general mechanism for the deleterious effect of introgressed ancestry.

## 1 Introduction

DNA introgressed from one population’s gene pool into another’s via hybridization is usually deleterious to the recipient population. Why this should be so has been a central question in evolutionary biology since the modern synthesis (Dobzhansky 1937; Muller 1942; Coyne and Orr 2004; Moran et al. 2021). More recently, patterns of introgression in population genomic data have suggested that the deleterious effect of introgressed ancestry is often spread across many loci genome-wide. For example, it has been estimated that Neanderthal variants have been deleterious to humans at thousands of loci across our genome (Juric et al. 2016), with similar estimates obtained for other species as well (e.g., Aeschbacher et al. 2017).

The classical theory for why introgressed ancestry is deleterious, due primarily to Dobzhansky and Muller, invokes negative fitness epistasis between variants that accumulated in the two populations during their period of divergence (Dobzhansky 1937; Muller 1942). These variants are neutral or beneficial against the genomic backgrounds of the populations in which they substituted, but, being untested against each other, can be incompatible, with these incompatibilities exposed in hybrids and their descendants after secondary contact between the populations (Orr 1995; Turelli and Orr 2000). Examples of Dobzhansky– Muller incompatibilities have been identified in many hybrid systems (e.g., Presgraves 2003; Powell et al. 2020).

A second kind of ‘incompatibility’ theory posits that introgressed variants, which are adapted to the donor population’s ecology, might be maladapted to aspects of the ecology of the recipient population (Mayr 1942; Schluter 2009). Barriers to gene flow due to ecological factors—both biotic and abiotic—have also been identified in a wide range of species (reviewed in Schluter 2000, 2009; Nosil 2012; Thompson et al. 2023).

Most recently, it has been noted that introgressed ancestry can be deleterious genome-wide if the donor species has a smaller effective population size—and therefore a greater genetic load—than the recipient species (Harris and Nielsen 2016; Juric et al. 2016). This scenario has been argued to pertain to Neanderthal–human introgression, given Neanderthals’ low effective population size relative to that of modern humans (Prüfer et al. 2014; Li et al. 2024).

Here, we explore an additional, potentially more general mechanism for the deleterious effect of introgressed DNA. We show that the regular operation of natural selection on complex traits—by which we mean stabilizing selection around trait optima—generates selection against the minor-parent ancestry in admixed populations. This is true even if the trait optima were the same for the parent populations (in contrast to theories of ecological incompatibilities) and if the parent populations were demographically identical (in contrast to load-based explanations). Since stabilizing selection on quantitative traits is ubiquitous, our results suggest that stabilizing selection could be a very general mechanism for selection against introgressed ancestry and the formation of reproductive barriers between species.

In parallel work to ours, Ragsdale (2024) has independently identified this mechanism of selection against introgression and demonstrated its effectiveness in demographically explicit genomic simulations of human–Neanderthal admixture.

## 2 Results

### Model

We study a simple model in which an ancestral population splits into two subpopulations, which diverge genetically during a period of isolation from one another and then combine in some proportions to form an admixed population. We are interested in the consequences of stabilizing selection on a quantitative trait for how the two ancestries’ proportions are expected to change over time in the admixed population.

We consider a single, highly polygenic, normally distributed, additive quantitative trait. Genetic variation in the trait is underlain by *L* ≫ 1 biallelic loci which recombine freely with one another. We consider autosomal loci only, postponing discussion of sex linkage to future work. An individual’s trait value, *Y*, is determined by

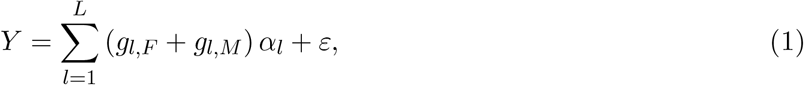

with *g*_*l,F*/*M*_ = −1/2 if the individual inherited the trait-decreasing allele at locus *l* from their father/mother, and *g*_*l,F*/*M*_ = +1/2 if they instead inherited the trait-increasing allele from their father/mother. *α*_*l*_ *>* 0 is the effect-size at the locus; in our calculations below, we assume this to be equal across loci (to a common value *α*). *ε* ∼ *N* (0, *V*_*E*_) is the environmental contribution to the trait.

We assume that the trait is under stabilizing selection and mark the trait optimum as 0; the relative fitness of an individual with trait value *Y* is given by

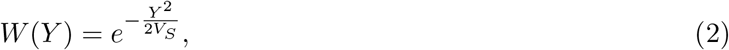

with *V*_*S*_ parameterizing the strength of stabilizing selection (with smaller values corresponding to stronger selection). For our analytical results, we assume that selection is weak, in the sense that the reduction of mean fitness due to stabilizing selection is small (1 − 𝔼[*W* (*Y*)] ≪ 1).

We consider the simplest demographic scenario: the ancestral population recently split into two populations, 1 and 2, of identical and constant size *N*. When we say that the ancestral population split ‘recently’, we mean that the time *T* since the split satisfies (i) *T* ≪ 2*N*_*e*_, where *N*_*e*_ is the effective population size, and (ii) *T* ≪ 1/*µ*, where *µ* is the mutation rate. These conditions ensure that the degree of divergence between the populations 1 and 2 satisfies *F*_*ST*_ ≪ 1 and that genetic variance in both populations is due almost entirely to loci that segregated in the ancestral population. (The assumption of recent divergence is important for our mathematical results below, but, as we argue in the Discussion, not for the conclusion that stabilizing selection disfavors the minor-parent ancestry in the admixed population.) After the period of divergence, the frequencies of the trait-increasing allele at locus *l* in populations 1 and 2 are 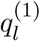 and 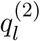 respectively.

The mean value of the trait in population *i* at the end of the period of divergence is 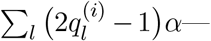 owing to stabilizing selection, these will equal the optimal value of the trait, 0, in expectation. Because the populations are demographically identical, their genetic variances 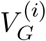 will also be equal in expectation (to a common value that we label *V*_*G*_), as will their genic variances 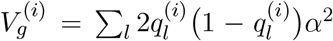 (to a common value that we label *V*_*g*_). (The genetic variance takes into account linkage disequilibria among loci, while the genic variance does not.)

In generation 0, a new, admixed population is formed, of the same size *N* and with a fraction 1 − *θ* of its ancestors deriving from population 1 and *θ* deriving from population 2. We assume that *θ <* 1/2, so that population 2 contributes the minor-parent ancestry; we will sometimes refer to population-2 ancestry in the admixed population as ‘introgressed’.

Generations are non-overlapping and mating is assumed to be random throughout (including with respect to ancestry in the admixed population).

### Intuition from the first two generations after admixture

Before presenting our general analytical approach, we first develop some basic intuition for the effect of stabilizing selection on the two ancestry proportions in the admixed population, by concentrating on the first two generations after admixture and assuming that the initial fraction of population-2 ancestry is very small.

Because population 1 is the major ancestry in the admixed population, most population-2 parents in generation 0 mate with population-1 parents. Therefore, most population-2 ancestry in generation 1 is contained in F1 hybrids. Similarly, most population-1 parents in generation 0 mate with fellow population-1 parents, so that, in generation 1, most population-1 ancestry is contained in individuals of pure population-1 ancestry (Fig. 1). If, in generation 1, F1 hybrids have lower fitness on average than individuals of pure population-1 ancestry, selection would reduce the fraction of population-2 ancestry between generations 1 and 2.

**Figure 1:**
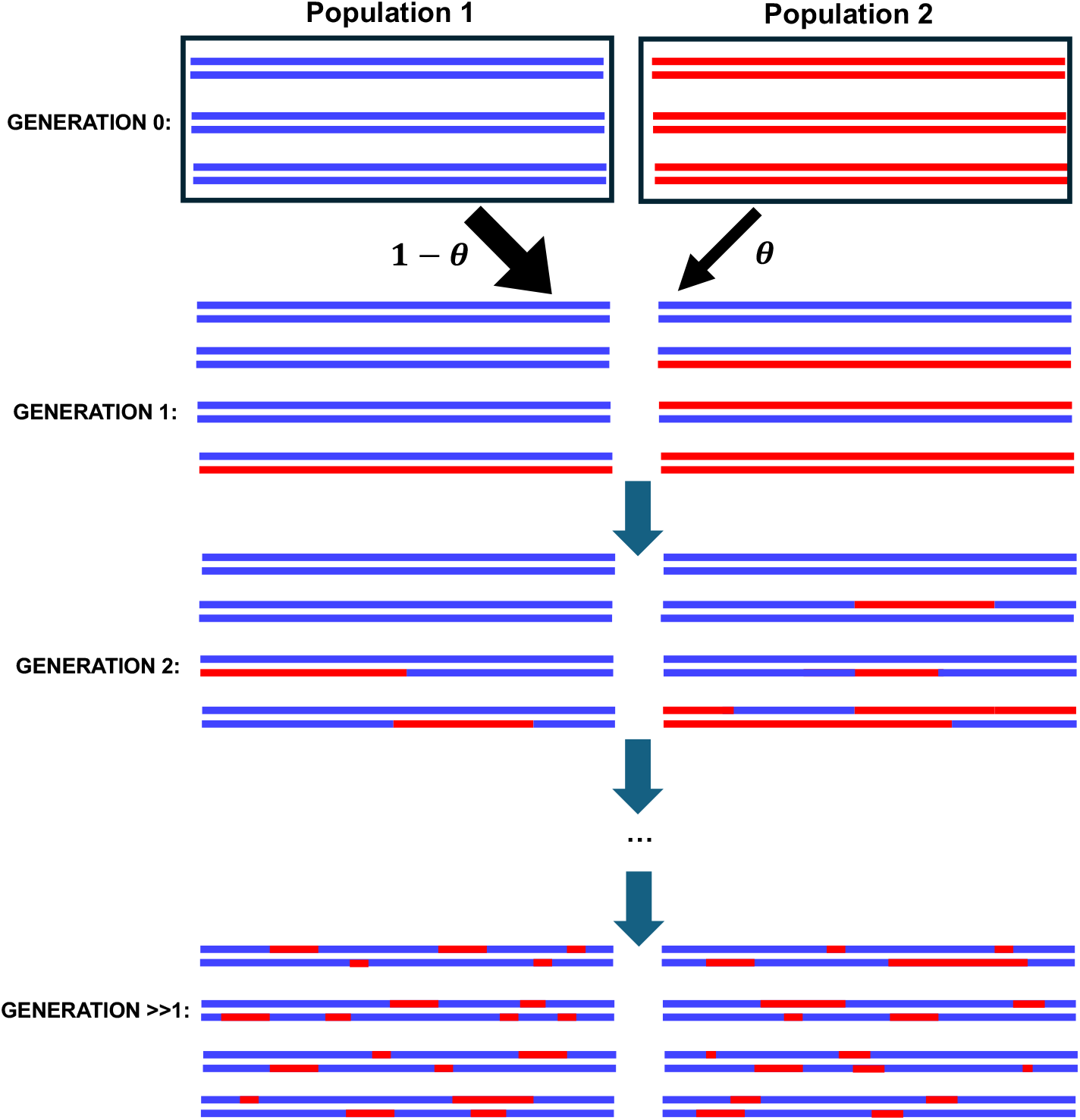
Schematic of the genomes of individuals in the first few generations after admixture.

However, this is not the case. Each F1 hybrid is formed from a gamete contributed by a population-1 parent and a population-2 parent, and the hybrid’s genetic value is the sum of the genetic values of these two gametes. Since, in each of the source populations, allele frequencies had co-adapted genome-wide to ensure a mean trait value of 0 (the optimum) and a genetic variance of *V*_*G*_, gametes derived from either source population have mean genetic value 0 and genetic variance *V*_*G*_/2. The mean and variance of the genetic values of F1 hybrids are therefore 0 and *V*_*G*_, the same as individuals of pure population-1 ancestry (as well as individuals of pure population-2 ancestry). Therefore, F1 hybrids are equally as fit, on average, as individuals of pure ancestry, and so there is no change in ancestry proportions expected between generations 1 and 2.

We now focus on selection in generation 2. Because most population-2 ancestry in generation 1 is contained in F1 hybrids, and because most of these hybrids mate with individuals of pure population-1 ancestry, most of the population-2 ancestry in generation 2 is contained in backcross hybrid offspring (Fig. 1). As in generation 1, most of the population-1 ancestry in generation 2 is in individuals of pure population-1 ancestry, whose genetic mean and variance are 0 and *V*_*G*_ respectively. If the backcross hybrid offspring are less fit than individuals of pure ancestry, then selection will reduce the fraction of population-2 ancestry between generations 2 and 3. Is this the case?

For a trait under stabilizing selection within a population, the genome-wide mean genetic value is held close to the optimal value. However, the mean genetic value of a given portion of the genome is free to drift away from the optimal value, as long as the complementary portion of the genome adjusts to compensate for this drift, keeping the genome-wide mean at the optimum (see Muralidhar, in prep., for an interesting application of this logic to sex chromosome turnover). Importantly, this drift in the mean genetic value of a given portion of the genome is independent in separate populations, so that the mean genetic value of the portion derived from one population together with the complementary portion derived from another population will not generally have a mean genetic value equal to the optimum. Moreover, the average distance from the optimum of the overall mean genetic value of this mixed genome will tend to be larger, the more genetically divergent the two populations are.

We now apply this reasoning to the backcross hybrids in generation 2. Each is formed from a gamete of pure population-1 ancestry and a recombined gamete from their F1 parent. Because of the randomness of recombination and segregation, these recombined gametes each have a different partition of the genome into a half from population 1 and a half from population 2 (Fig. 2). Because each half derives from a different ancestry, per the reasoning above, every partition corresponds to a different deviation of the mean genetic value from the optimum of 0. These deviations can be positive or negative, and so the mean genetic value across all partitions—and thus all gametes produced by F1 parents—is still 0, but the genetic values show greater variation around this mean than the genetic values of gametes of pure ancestry.

**Figure 2:**
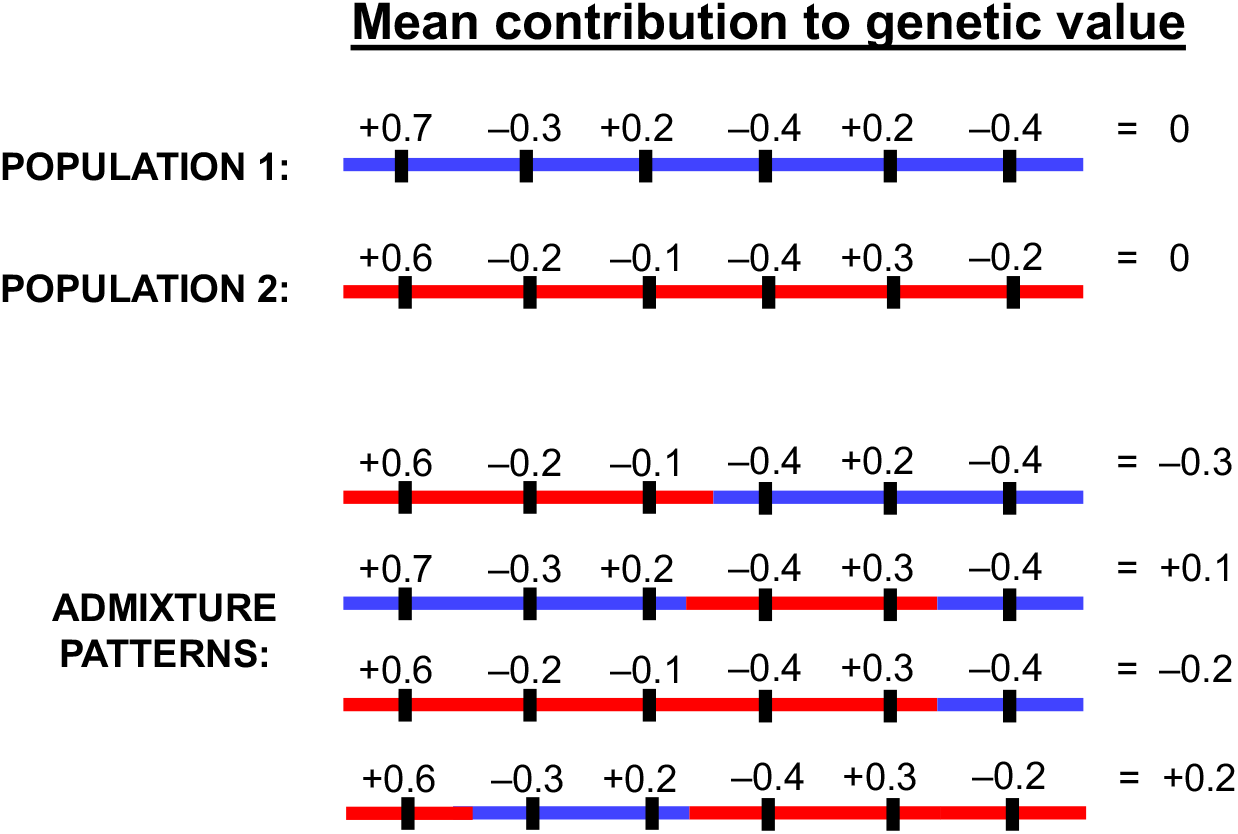
Admixture disrupts the within-population co-adaptation of allele frequencies across the genome. Each causal locus has a mean effect on the trait that depends on its alleles’ effect sizes (which are the same for the two populations) and frequencies (which may differ between the two populations). Within each parent population, allele frequencies at different loci have co-adapted under stabilizing selection to keep the mean genetic contribution of the entire genome close to zero (the optimal value). However, when a portion of the genome is sampled from population 1 and the complementary portion is sampled from population 2, allele frequencies within the two portions need not be co-adapted, and the mean genetic value of the resulting genome can deviate from zero.

Therefore, while the mean genetic value among backcross offspring in generation 2 is 0, these offspring show greater variance in their genetic values than offspring of pure ancestry do (Slatkin and Lande 1994; Barton 2001; Schneemann et al. 2020; Moran et al. 2021). Under stabilizing selection, fitness declines with the distance of the trait from the optimum, so that a group of individuals with greater trait variation has lower average fitness than a group of individuals with the same mean trait value but lower variance. As a result, backcross offspring in generation 2 are, as a group, less fit on average than offspring of pure ancestry. Therefore, population-2 ancestry declines from generation 2 to 3.

The intuition above suggests that selection against population-2 ancestry, beginning in the second generation after admixture, will be stronger if (i) populations 1 and 2 are more genetically diverged, so that, when the genome is partitioned into regions of unlike ancestry, these regions are less co-adapted, and (ii) the initial ancestry fraction deriving from population 2 (*θ*) is smaller, so that a greater fraction of population-2 ancestry resides in hybrids. In the next section, we will derive a general expression for the change in ancestry fractions in each generation after admixture.

### Genome-wide change in ancestry

In order to quantify the change in ancestry per generation in the admixed population, we will generalize the intuition obtained above. We will first derive an expression for the increase in the genetic variance of gametes produced by individuals of a given ancestry fraction (where an individual’s ‘ancestry fraction’ is defined as the fraction of their genome that derives from source population 2). We will then use this expression to calculate the mean fitness of offspring as a function of the ancestry fractions of their parents. Finally, we will substitute these mean fitnesses, along with the distribution of ancestry in each generation, into a breeder’s equation to calculate the expected change in the mean ancestry fraction each generation after admixture.

#### The genetic variance of gametes produced by hybrid parents

Consider the gametes produced by individuals with ancestry fraction *p*, and focus on a trait-affecting (or ‘causal’) locus, indexed by *l*. What is the contribution of this locus to genetic variance in the gametes produced by individuals with ancestry fraction *p*? We calculate this contribution using the law of total variance:

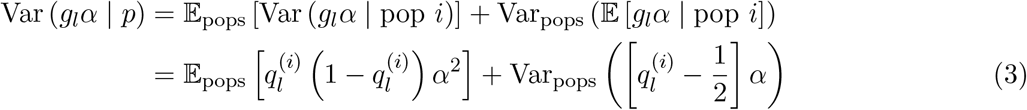

where *α* is the effect size (assumed to be the same across loci), *g*_*l*_ is the genotype at locus *l* in the gamete (−1/2 if the trait-decreasing allele; +1/2 if the trait-increasing allele), and 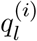 is the frequency of the trait-increasing allele in source population *i*. The subscript ‘pops’ indicates that the expectation and variance are taken over the distribution of ancestries of the allele at the locus in gametes produced by individuals with ancestry fraction *p*. This distribution is Bernoulli; because of our assumption of weak selection, with probability 1 − *p* the allele derives from population 1 and with probability *p* it derives instead from population 2. Therefore, the first term in Eq. (3) is a weighted sum of the contributions of locus *l* to genetic variance in gametes in populations 1 and 2:

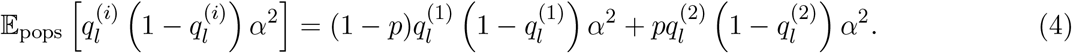

The second term is where the additional genetic variance in the admixed gametes comes from:

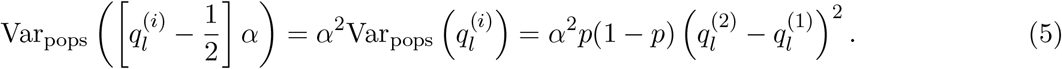

From our assumptions that selection is weak and that there are many unlinked causal loci, and ignoring signed linkage disequilibrium among trait-increasing alleles, we can sum the contribution in Eq. (5) across loci *l* = 1, 2, …, *L* to find the overall genetic variance of gametes produced by individuals with ancestry fraction *p*:

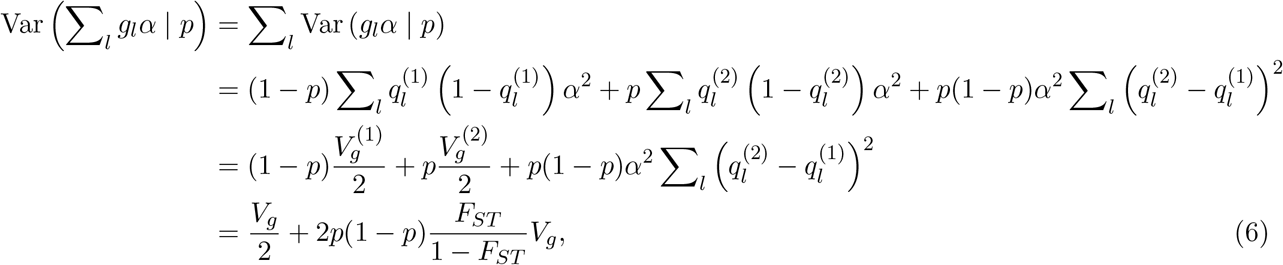

where 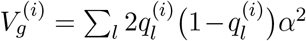 is the genic variance for the trait in population *i*, and the last line follows from our assumption that these genic variances are equal (to *V*_*g*_) in the two source populations and the fact that, as we show in the Methods, 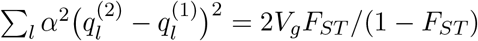, with *F*_*ST*_ measuring the degree of genetic divergence between the two source populations. In the Methods, we also derive Eq. (6) more formally by considering all possible sets of loci constituting a fraction *p* of the haploid genome. We also show in the Methods that there is a small correction to the *F*_*ST*_ /(1 − *F*_*ST*_) term in Eq. (6) due to long-range signed linkage disequilibrium in the parent populations, which could arise, even under free recombination, because of the action of stabilizing selection itself (Bulmer 1971) or because of forces such as population structure.

We can see from Eq. (6) that the increase in the genetic variance of gametes is maximal for individuals with ancestry fraction *p* = 1/2, and goes to zero for individuals of pure population 1 or 2 ancestry. Moreover, the additional genetic variance in admixed gametes increases with the degree of genetic divergence *F*_*ST*_ of the two source populations, becoming very large if *F*_*ST*_ is close to 1.

#### The average fitness of offspring with given parental ancestry fractions

Each offspring derives from two gametes, one from their mother, whose ancestry fraction we label *p*_*M*_, and one from their father, whose ancestry fraction we label *p*_*F*_. The offspring’s ancestry fraction is (*p*_*F*_ + *p*_*M*_)/2 in expectation. Because mating is assumed to be random, the genetic variance among such offspring, and thus their average fitness, is the sum of the genetic variances of the gametes that produced them. These are given by Eq. (6), and so the genetic variance among offspring with parental ancestry fractions *p*_*F*_ and *p*_*M*_ is

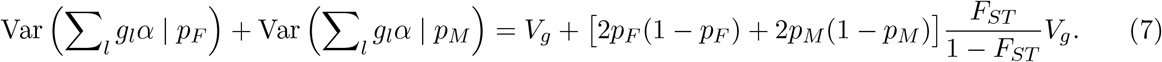

Under weak stabilizing selection, the mean relative fitness of a group of individuals whose phenotypic values are distributed normally around the optimum with variance *V*_*P*_ is approximately 1 − *V*_*P*_ /2*V*_*S*_. The mean relative fitness of these offspring, whose phenotypic variance is given by the genetic variance in Eq. (7) plus environmental variance *V*_*E*_, is therefore

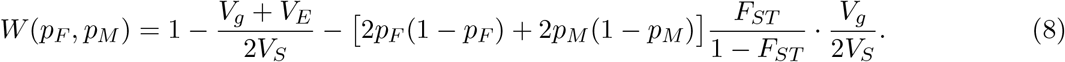

#### The change in ancestry across a generation

Eq. (8) allows us to calculate the expected change in ancestry fraction from one generation to the next. Let *Z*_*t*_ be the fraction of introgressed (population-2) ancestry among zygotes in generation *t*. We want to calculate Δ*Z*_*t*_ = *Z*_*t*+1_ − *Z*_*t*_. Let 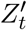 be the fraction of introgressed ancestry among adults after selection has acted in generation *t*, and let the distribution of *p*_*F*_ and of *p*_*M*_ among the gametes produced by these adults be *f*_*t*_(*p*) (which is the same for males and females). These various quantities are related by

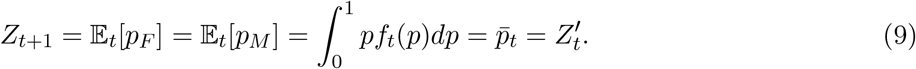

The ancestry fraction *Z*_*t*+2_ among zygotes in generation *t* + 2 equals the ancestry fraction 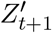 among adults after selection in generation *t* + 1, which in turn is given by

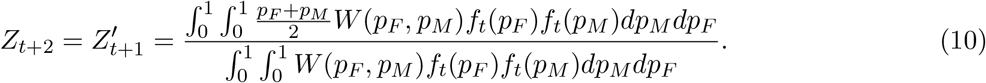

Eq. (10) makes use of the assumption of random mating. In the Methods, we use Eq. (10) to calculate that, to first order in *V*_*g*_/*V*_*S*_ and *V*_*E*_/*V*_*S*_,

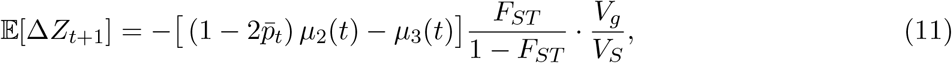

where 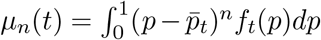 is the *n*-th central moment of *f*_*t*_(*p*). Thus, the change in mean ancestry fraction between generations *t* + 1 and *t* + 2 depends on the second and third central moments of the distribution *f*_*t*_(*p*) of ancestry fractions *p*_*F*_ and *p*_*M*_ among gametes (and thus adult individuals) in generation *t*.

As long as the number of generations *t* since admixture is not too large relative to (the log of) the population size, an individual in generation *t* descends from 2^*t*^ distinct ancestors in generation 0. Under weak selection, these ancestors are ‘sampled’ with uniform probability from the generation-0 population, so that the probability that a given ancestor is from population 2 is *θ*. Therefore, the number of population-2 ancestors *k* among the 2^*t*^ total ancestors of a randomly chosen individual in generation *t* is distributed binomially with 2^*t*^ trials and probability *θ*; the mean, second central moment, and third central moment of *k* are 2^*t*^*θ*, 2^*t*^*θ*(1 − *θ*), and 2^*t*^*θ*(1 − *θ*)(1 − 2*θ*), respectively. Because there are many loci, all unlinked, and selection is weak, the individual receives a fraction 1/2^*t*^ of their genome from each ancestor, and therefore a fraction *k*/2^*t*^ from population 2. So the distribution *f*_*t*_(*p*) of the fraction of individuals’ genomes that derives from population 2, *p*_*t*_ = *k*/2^*t*^, has moments

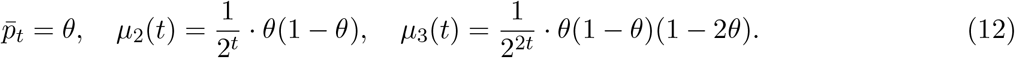

Selection induces correction terms to these expressions of the order of *V*_*g*_/*V*_*S*_, which we can ignore in the limit of weak selection. When considering the effects of strong stabilizing selection, these higher-order terms would need to be taken into account.

Substituting these expressions for the moments of *f*_*t*_(*p*) into Eq. (11), we find that

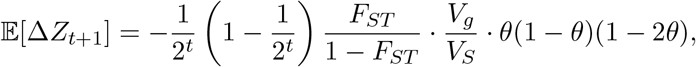

or, adjusting time indices,

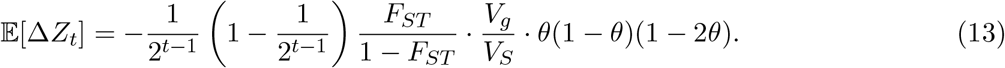

Eq. (13) is the expected change in the fraction of population-2 ancestry in the admixed population between generations *t* and *t* + 1. It is zero when *t* = 1, validating our argument that there is no selection in favor of either ancestry in the first generation after admixture because F1 hybrids are as fit, on average, as individuals of either pure ancestry. However, when *θ <* 1/2, Eq. (13) is negative in each generation *t* ≥ 2 after admixture, r evealing)that the minor ancestry will be selected against over time. Furthermore, because the term 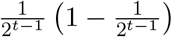 shrinks as *t* grows, Eq. (13) shows that the expected decline of the minor ancestry is fastest in the first few generations after admixture, but gradually slows down in later generations. This effect is observed in our simulations (Fig. 3A), and is a common feature of models of selection against introgressed ancestry (Harris and Nielsen 2016; Veller et al. 2023). Additionally, the rate at which introgressed ancestry is expected to be purged (and the eventual amount of purging) is greater if the degree of divergence *F*_*ST*_ between the source populations is larger (since 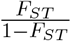 is increasing in *F*_*ST*_) and if stabilizing selection is stronger (smaller *V*_*S*_).

**Figure 3:**
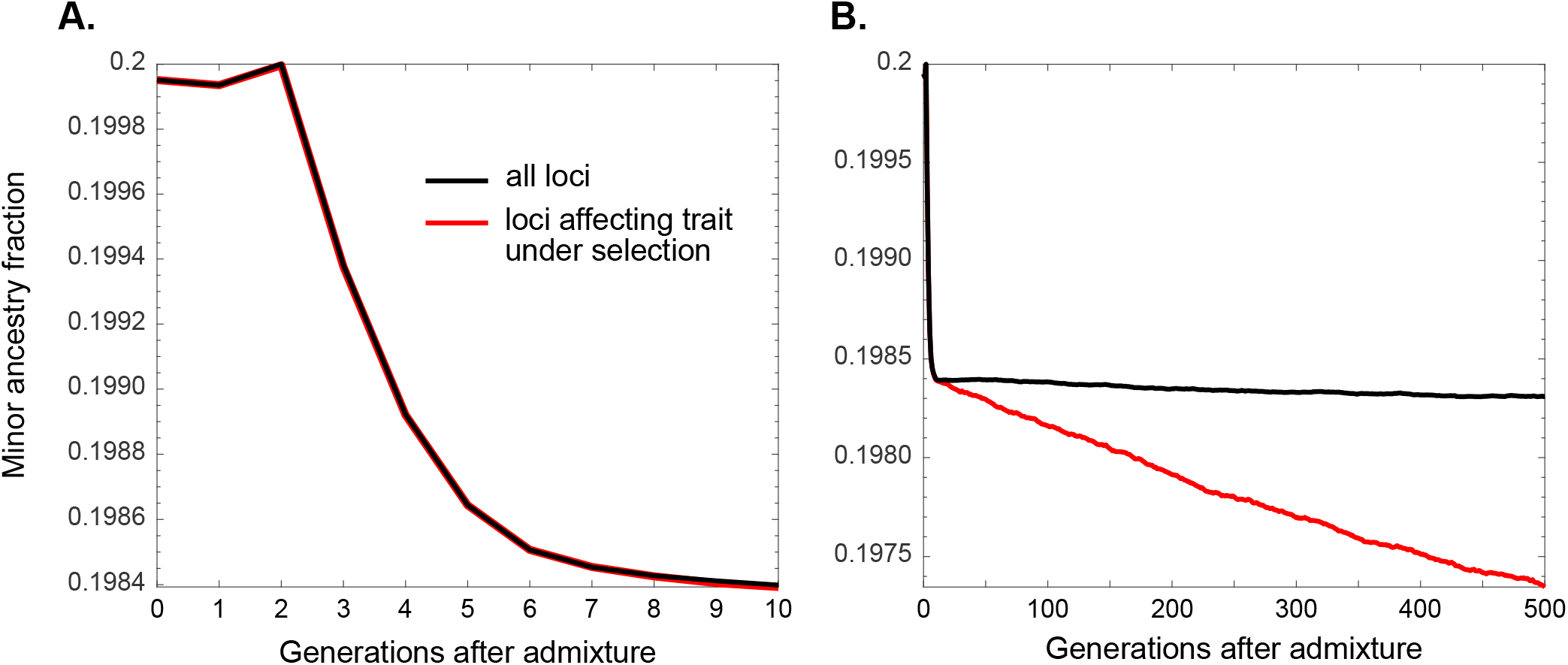
Stabilizing selection generates selection against the minor ancestry in an admixed population. Average minor-parent ancestry over time, with free recombination between all loci. (**A**.) In the early generations after admixture, selection against the minor ancestry is rapid, and approximately equal at loci that do and do not affect the trait under selection (because recombination has not yet fully decoupled ancestry at these loci). (**B**.) In later generations after the admixture event, there is no selection against the minor ancestry at loci that do not affect the trait under selection, because recombination has fully decoupled the ancestry at these loci from the ancestry at causal loci. However, there is slow long-term selection against the minor ancestry at causal loci. In the simulations displayed here, the strength of selection is *V*_*S*_/*V*_*g*_ = 8, the initial minor-parent ancestry fraction is *θ* = 0.2, and the degree of divergence of the source populations is *F*_*ST*_ = 0.2. Trajectories are averaged across 1,000 replicate trials. Full details of the simulations can be found in the Methods.

Summing the per-generation changes in Eq. (13) across the *t* generations since admixture yields:

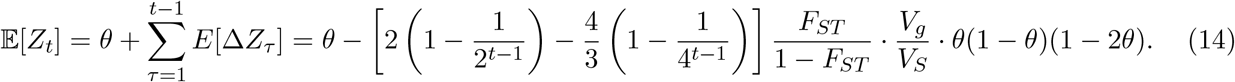

For *t* ≫ 1, this converges to

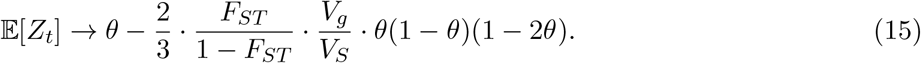

Eq. (15) can be interpreted as the expected introgressed fraction, many generations after admixture, at loci that do not affect the trait. Because of our assumption of no linkage, when *t* ≫ log_2_(*L*) (where *L* is the number of loci that affect the trait), the ancestry at any locus that does not affect the trait has become decoupled from the ancestry at loci that do affect the trait (‘causal’ loci/alleles), and therefore also uncorrelated with trait variance. So, if the allele was introgressed, it eventually becomes unaffected by stabilizing selection despite its initial association with causal introgressed alleles, conditional on its having survived this initial association (Fig. 3B).

By this interpretation, Eq. (15) provides the ‘gene-flow factor’ (Bengtsson 1985; Barton and Bengtsson 1986) associated with stabilizing selection’s effect on introgressed ancestry. Among the fraction *m* = *θ* of neutral alleles that migrate into the recipient population in the introgression event, the proportion *m*_*e*_ that survive (and thus ‘effectively migrate’) in the recipient population is given by Eq. (15). The gene-flow factor is

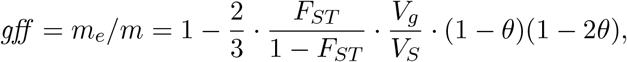

or, when *θ* ≪ 1,

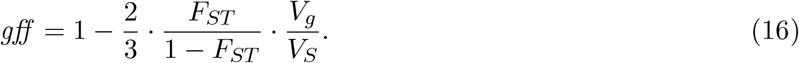

Early after admixture, ancestry dynamics are similar at causal and neutral loci, because neutral introgressed alleles retain their association with some of the causal alleles with which they introgressed (Fig. 3A). Once neutral and causal introgressed alleles have become decoupled, such that neutral alleles have become unaffected by stabilizing selection, ancestry at causal loci remains correlated with trait variance, and so stabilizing selection affects ancestry dynamics at these loci even in the long run. In the next section, we repeat the calculations above but consider ancestry fractions only at causal loci.

### Long-term change in ancestry at causal loci

In the first few generations after admixture, the fate of introgressed ancestry at a given causal locus is largely determined by the many other causal loci in the long ancestry blocks in which the locus is contained, rather than by the individual effects of the alleles at the locus itself. However, when *t* ≫ log_2_(*L*), where *L* is the number of causal loci for the trait, recombination will have broken up the initial long ancestry blocks such that they now contain at most one causal locus each. In this case, consider Eq. (13) applied to these *L* causal loci, and in particular, consider the moments of the distribution *f*_*t*_ of *p*_*t*_, the fraction of introgressed ancestry at causal loci in gametes (and thus individuals) in generation *t* ≫ log_2_(*L*). Assuming that the source populations are sufficiently large relative to *L*, the alleles at the *L* causal loci in a generation-*t* gamete all derive from distinct ancestors in the original admixed population; the number *k* that derive from introgressing ancestors is therefore, under weak selection, binomially distributed with *L* trials and probability *θ*; the mean, second central moment, and third central moment of this binomial distribution are *Lθ, Lθ*(1 − *θ*), and *Lθ*(1 − *θ*)(1 − 2*θ*). The fraction of introgressed ancestry at causal loci in the gamete is *p* = *k*/*L*; the moments of its distribution *f*_*t*_(*p*) are then

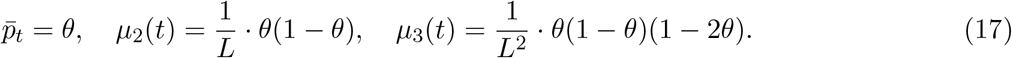

Unlike for the case of neutral loci (Eq. 12), the moments above do not depend on time *t*, because beyond *t* ≫ log_2_(*L*), the number of ancestors of the alleles at the *L* causal loci in a gamete does not grow with *t*.

Substituting these moments into Eq. (11), we find that, at causal loci, many generations after admix-ture,

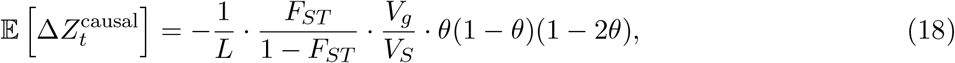

where we have ignored terms of order 1/*L*^2^.

Thus, the initial rapid decline in introgressed ancestry at causal and neutral loci is followed by a slower, perpetual decline at causal loci (Fig. 3B).

#### Accounting for the linkage disequilibrium that stabilizing selection generates

Stabilizing selection generates positive linkage disequilibria genome-wide between alleles with opposite directional effects on the trait (Bulmer 1971), partially masking the individual effects of these alleles and therefore slowing down their frequency dynamics. Standard formulae for the speed of allele-frequency dynamics under stabilizing selection do not take into account this slowdown due to the LD that stabilizing selection necessarily generates, but Negm and Veller (2024) show that this can be achieved simply by defining an ‘effective’ effect size of the alleles at each locus as a function of the overall amount of LD generated by stabilizing selection and the locus’s recombination relations with loci elsewhere in the genome.

In the case where all loci are unlinked and have equal effect sizes, the ‘effective’ effect size of an allele is reduced from its true effect size *α* by a factor 1 − *δ*, where 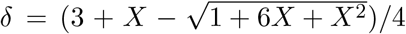, with *X* = (*V*_*S*_ + *V*_*E*_)/*V*_*g*_. The expected change in frequency of the allele, which under standard formulae is proportional to *α*^2^ (reflected in Eq. 18 in the numerator term *V*_*g*_), is therefore slowed by a factor (1 − *δ*)^2^.

Additionally, Eq. (18) does not take into account the effect of background phenotypic variance on the selection gradient; doing so, accounting for the effect of linkage disequilibrium on this phenotypic variance, introduces a further slowdown factor 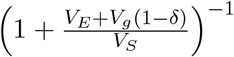 (Negm and Veller 2024), so that a more accurate prediction of the change in ancestry fraction at causal sites many generations after admixture is

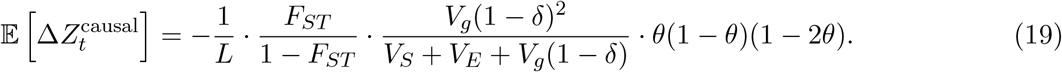

Eq. (19) predicts the long-term rate of purging of introgressed ancestry at causal loci in our simulations more accurately than Eq. (18) does. For example, in the simulations displayed in Fig. 3, the observed slope of the trajectory of the introgressed fraction at causal loci between generations 50 and 500 is −2.15×10^−6^; Eq. (19) predicts a value of −2.22 × 10^−6^, while Eq. (18) predicts a value of −3.00 × 10^−6^. The greater accuracy of Eq. (19) carries over to other parameter configurations as well (Fig. S1).

### The effects of linkage

Thus far, we have assumed free recombination between all loci, causal and neutral. Simulations with realistic linkage maps reveal that linkage can have a substantial effect on the fate of introgressed ancestry under stabilizing selection (Fig. 4).

**Figure 4:**
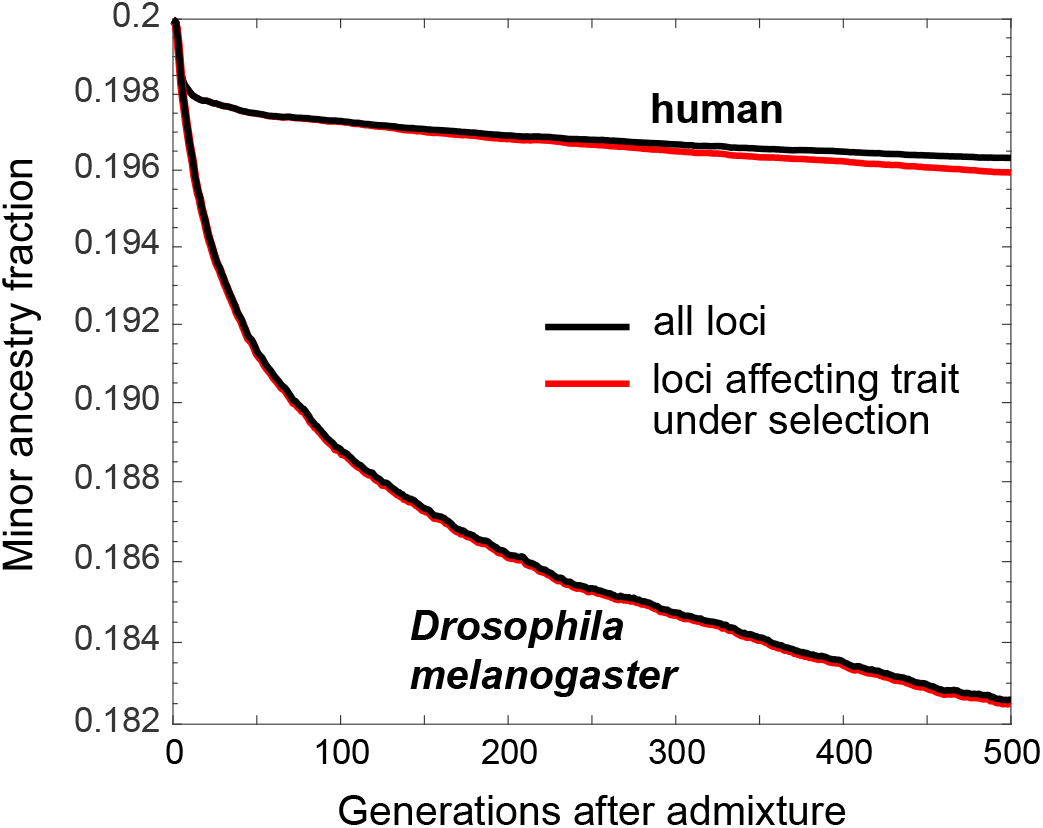
Linkage increases the rate of purging of the minor ancestry under stabilizing selection. Shown are the average minor-parent ancestry fractions at causal and neutral loci, under the recombination processes of humans and *Drosophila melanogaster*. All other parameters are as in Fig. 3, and trajectories are averaged across 1,000 replicate trials. Full details of the simulations can be found in the Methods.

For simulations carried out with the human recombination map (Kong et al. 2010) and the linkage map of *Drosophila melanogaster* (Comeron et al. 2012), the rate of purging of introgressed ancestry at causal sites is faster than under free recombination, and the introgressed fraction at neutral sites hews more closely to that at causal loci (Figs. 3,4). These observations reflect two effects of linkage on selection against introgressed ancestry (Veller et al. 2023): (i) it keeps introgressed alleles at causal loci together for a longer time, strengthening selection against them, and (ii) it keeps neutral introgressed alleles together with their causal counterparts for longer, increasing the probability that the neutral alleles will be eliminated together with the causal alleles. These effects are particularly noticeable in the case of *Drosophila melanogaster* (Fig. 4), a low-recombination species with only two major autosomes and no crossing over in males.

## 3 Discussion

We have shown, via analytical calculations and whole-genome simulations, that stabilizing selection on quantitative traits can generate selection against the minor-parent ancestry in admixed populations. This suggests a potentially very general mechanism, identified also in parallel work by Ragsdale (2024), for selection against introgressed DNA and, more broadly, for the formation of reproductive barriers between populations and species. However, since we have focused here on the simplest possible model—with two recently diverged and demographically identical populations under the same regime of stabilizing selection on a single trait—we see our results primarily as demonstrating proof of principle. Below, we discuss several directions in which our results might be generalized theoretically and applied empirically.

### Linkage and linkage disequilibrium

Perhaps the most pertinent effect that our analytical calculations have ignored is the role of the recombination map in modulating the rate at which minor-parent ancestry is purged, though our simulations have revealed that this role can be substantial (Fig. 4).

In the alternative, simpler case of additive selection against introgressed ancestry (as might occur, for example, if the minor-parent population suffers a greater genetic load), it has been shown that the effect of linkage on the fate of introgressed ancestry can be captured analytically as a function of certain metrics of the aggregate recombination process (Veller et al. 2019) which control ancestry variance in the admixed population (Veller et al. 2023). In future work, we plan to extend our calculations above in the same line as for additive selection against introgressed ancestry, to characterize the effect of linkage on the fate of introgressed ancestry under the stabilizing selection mechanism that we have identified.

Though our calculations have also largely ignored the effects of linkage disequilibrium, we have shown that the LD that is expected to build up under stabilizing selection (Bulmer 1971) can be taken into account by including a ‘slowdown’ factor (Negm and Veller 2024) in the equation describing the long-term rate of purging of minor-parent ancestry at causal loci. Moreover, we have shown that the effect of LD present in the source populations, owing perhaps to selection, population stratification, or other forces, on the genetic variance of hybrids in the admixed population can also be calculated (Methods). However, a more complete analysis of the effects of LD on the purging of introgressed ancestry under stabilizing selection is required, and will be the subject of future work.

### The degree of divergence of the parent populations

Our calculations rely to some extent on an assumption of recent divergence of the source populations, such that most of the genetic variation for the trait under selection is due to polymorphic loci that are shared between the two populations. However, stabilizing selection will itself accelerate independent turnover of the genetic variation in the two populations (Yair and Coop 2022), and admixture of more anciently diverged source populations is in any case of obvious empirical interest. Like for weakly divergent source populations, the conclusion that stabilizing selection disfavors the minor ancestry in admixed populations holds also in the case of highly divergent source populations, with an even more forceful logic.

This is clearest for fixed differences between the populations. Consider a locus at which a trait-increasing allele is fixed in one population and a (relatively) trait-decreasing allele is fixed in the other population. Upon hybridization, the locus becomes polymorphic in the admixed population, and the allele contributed by the minor-parent ancestry is, by definition, the rare allele at the locus. Since stabilizing selection selects against rare alleles (Robertson 1956), it selects against the minor-parent ancestry in this case.

Similarly, consider a locus that is fixed in one population but polymorphic in the other. Under our assumption that the source populations are demographically identical, half of such loci will be polymorphisms private to population 1 and the other half will be private to population 2; the frequency spectra of these two sets of loci will also be the same. Upon hybridization, the alleles that were private to the minor-parent population always become the rare alleles at their loci in the admixed population, and, moreover, will be rarer on average than the alleles that were private to the major-parent population; they are therefore selected against more strongly on average than the alleles that were private to the major-parent population, and so, again, stabilizing selection disfavors the minor-parent ancestry.

In the Methods, we extend our calculations to cover the case of highly divergent source populations, between which there are some number of fixed differences at trait-affecting loci, and within which the genetic variation is contributed by private polymorphisms. We show that, sensibly, selection against the minor ancestry is stronger in this case than for weakly divergent populations with shared genetic variation.

### The number of traits under stabilizing selection

We have considered the effect of weak stabilizing selection on a single trait. However, many quantitative traits are likely under stabilizing selection (Sanjak et al. 2018; Simons et al. 2018; Sella and Barton 2019), and certain traits will sometimes be under strong stabilizing selection (e.g., de Villemereuil et al. 2020). How stabilizing selection on many traits will affect the dynamics that we have described here will depend on the number of traits under selection, the strengths of selection on them, and the degree and nature of pleiotropy of the alleles underlying variation in these traits.

Most obviously, selection on many traits will greatly increase the number of ‘causal’ loci that are subject to direct, rather than correlated, selection against the minor-parent ancestry. Moreover, the increased density of these causal loci throughout the genome will reduce the average recombination distance of neutral loci to their nearest causal loci, increasing the degree to which the dynamics at neutral loci reflect the correlated effects of direct selection against minor-parent ancestry at the causal loci. The ultimate result will be a greater purging of minor-parent ancestry genome-wide.

### Interactions with other mechanisms of selection against introgressed ancestry

When divergent populations come into secondary contact, several kinds of selective forces can disfavor introgression of DNA from one population into the other, including Dobzhansky–Muller incompatibilities (DMIs), environmental incompatibilities, and differences in mutation load between the parent populations (Moran et al. 2021). What distinguishes stabilizing selection from these other mechanisms, and how does it interact with them?

The stabilizing selection mechanism is distinct from ecological incompatibilities, since, as we have shown, it operates even when the admixing populations have identical ecological contexts, reflected in identical trait optima. Ecological incompatibilities can be incorporated into our model by allowing the trait optima to differ between the source populations, with the admixed population subject to one of the trait optima (or perhaps a new trait optimum).

Stabilizing selection is also mechanistically distinct from load-based arguments that rely on differences in the effective population sizes of the admixing populations, since, as we have shown, stabilizing selection will select against the minor-parent ancestry even when the source populations are demographically identical. It is, however, interesting to consider the interaction between stabilizing selection and differences in the effective sizes of the admixing populations, since the effects of population size on allele-frequency dynamics under stabilizing selection are complex (Simons et al. 2014, 2018). In this regard, Ragsdale (2024) has carried out simulations of the stabilizing selection mechanism calibrated to the demography of human–Neanderthal admixture, including their unequal effective population sizes.

Among the alternative mechanisms for selection against introgressed DNA, stabilizing selection is perhaps conceptually closest to DMIs. We have argued that the effect of stabilizing selection in disfavoring the minor-parent ancestry can be understood in terms of a disruption of the co-adaptation of allele frequencies within populations under stabilizing selection. In each source population, the mean genetic value of a portion of the genome can drift freely, as long as the genetic value of the complementary portion adjusts so as to keep the genome-wide genetic value at the optimum. With this process operating independently in the two source populations, a portion of the genome from one population, though coadapted to the complementary portion from the same population because of stabilizing selection, need not be co-adapted to the complementary portion from the other population. This could be interpreted as a genetic incompatibility between the two portions of the genome when they derive from distinct populations.

Note, however, that a hybrid here will suffer reduced fitness in the case where it inherits the first genomic portion from population 1 and the second portion from population 2 and in the reciprocal case where it inherits the first portion from population 2 and the second portion from population 1. This is in contrast to the usual ‘asymmetric’ picture of DMIs (Muller 1942; Orr 1995), where, ‘if allele *A* from one species is incompatible with allele *B* from the other, alleles *a* and *b* must be compatible’ (Orr 1995, pg. 1811). A second (though related) difference between the stabilizing selection mechanism that we have elucidated here and DMIs is that it operates even when all of the loci involved harbor polymorphisms shared between the two populations. DMIs, in contrast, traditionally involve alleles that are derived in one population and absent in the other.

Stabilizing selection, as a mechanism for selection against introgressed DNA, is therefore conceptually distinct from these various other mechanisms, though it might interact with them in interesting ways. In general, it would be possible to model analytically, and to simulate, all of the mechanisms together in order to understand how their relative importance depends on the parameters of each mechanism.

### Empirical predictions and evidence

Since it is mechanistically distinct from ecological incompatibilities, load-based arguments, and DMIs, stabilizing selection differs from these mechanisms in its empirical predictions. A key difference between stabilizing selection and both ecological incompatibilities and load is that stabilizing selection selects specifically against the minor-parent ancestry in admixed populations. This property is shared with DMIs (Schumer et al. 2018; Moran et al. 2021), and it implies that stabilizing selection (and DMIs) can be empirically distinguished from ecological incompatibilities and load in cases of bi-directional gene flow between diverged populations, or in replicate hybrid populations where the source population that contributes the minor-parent ancestry is flipped.

Selection against minor-parent ancestry specifically has been observed in replicate admixed populations of swordtail fishes (Schumer et al. 2018). In the context of human–Neanderthal admixture, it has been known for some time that Neanderthal ancestry was likely broadly deleterious in humans (e.g., Sankararaman et al. 2014; Harris and Nielsen 2016; Juric et al. 2016), and more recent evidence suggests a deleterious effect of human ancestry in certain regions of the Neanderthal genome as well (Harris et al. 2023). This is consistent with the effect of stabilizing selection that we have identified here, and also with DMIs (as argued by Harris et al. 2023). However, evidence that human ancestry was deleterious in the Neanderthal genome in some of the same regions where Neanderthal ancestry was deleterious in the human genome (Harris et al. 2023)—that is, the existence of reciprocal introgression ‘deserts’—appears to be uniquely consistent with the stabilizing selection mechanism, since under the traditional model of DMIs, only one population would contribute the incompatible allele in each such region (Ragsdale 2024).

Observing the effect of the stabilizing selection mechanism in genomic data requires understanding how stabilizing selection on introgression affects the genetic architecture of traits and of introgressed ancestry. Thus, it would require understanding how the spread and length of introgressed genomic segments depend on the frequencies and effect sizes of genetic variants within and around these segments. This in turn requires an explicit model of the joint distribution of allele frequencies and effect sizes in the source populations and the admixed population. This joint distribution is shaped by stabilizing selection, as well as the relevant demographic scenarios, recombination maps, and other effects (Sella and Barton 2019). In addition, as discussed above, the stabilizing selection mechanism likely interacts with other forms of selection on introgressed ancestry, and one would need to also consider these other mechanisms not just as alternatives to stabilizing selection but also as mediating the effect of stabilizing selection.

Ultimately, a detailed understanding of the effect of stabilizing selection on introgression in natural populations will require whole-genome simulations under realistic demographic scenarios, patterns of linkage, strengths of selection, distributions of allelic frequencies and effect sizes, and patterns of pleiotropy across traits. A first look in this regard has been given by Ragsdale (2024).

### How important is stabilizing selection for the fate of introgressed ancestry?

For analytical tractability, we have made several strong assumptions in our calculations. These assumptions include: weak divergence of the source populations, weak stabilizing selection, a solitary quantitative trait under selection, and no linkage. Each of these assumptions has the effect of limiting the impact of stabilizing selection in purging the minor ancestry, and so, together, they restrict us to a parameter regime where stabilizing selection can have only a very limited effect on introgression. In realistic scenarios where all of these assumptions are relaxed, stabilizing selection may prove to be a quantitatively important force in selection against introgression and the formation of reproductive barriers between species.

## Acknowledgments

We are grateful to Graham Coop, Hunter Fraser, Priya Moorjani, Ben Moran, Pavitra Muralidhar, Sherif Negm, Dan Powell, Jonathan Pritchard, Aaron Ragsdale, Molly Schumer, Guy Sella, and Ken Thompson for helpful discussions. Funding was provided by a Branco Weiss Fellowship to CV.

## Methods

### Relating genetic variance and *F*_*ST*_

The frequency of the trait-increasing allele at locus *l* in source population *i* is 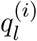. Write 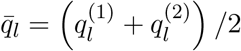. We assume that each locus *l* ∈ {1, 2, …, *L*} that affects the trait has the same effect size, *α*. Across these loci, the definition of *F*_*ST*_ that is relevant to our calculations is

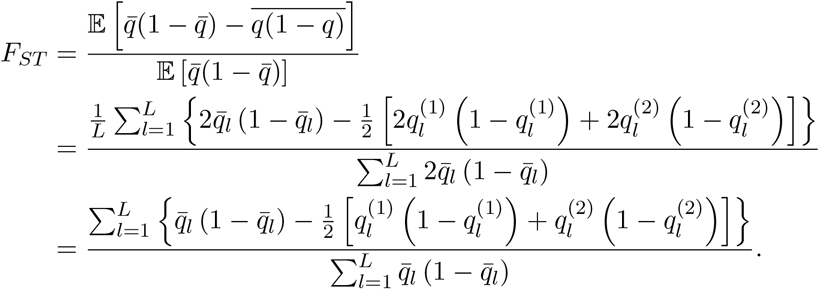

Rearranging and simplifying,

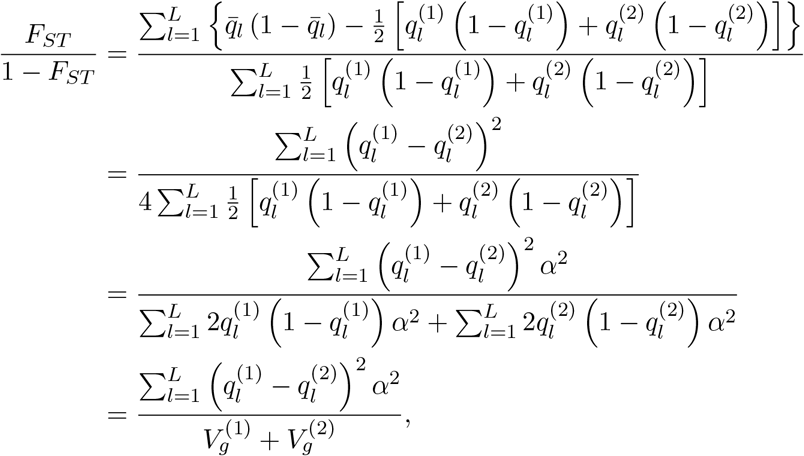

where 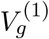 and 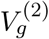 are the genic variances of populations 1 and 2. We assume these genic variances to be equal to a common value *V*_*g*_, so that

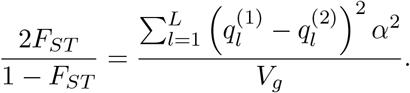

### Variance of the genetic values of gametes containing a certain fraction of introgressed ancestry

Consider a gamete in which a fraction *p* of the DNA derives from the introgressing population 2, with a fraction 1 − *p* deriving from population 1. Because of our assumptions of (i) weak selection, (ii) many causal loci, and (iii) no linkage, a number *I* = *pL* of the causal loci in the gamete derive from population 2 and, among the total number *L* of causal loci, any such set of *I* loci is equally likely. We want to calculate the variance of the genetic values among such gametes with a fraction *p* of introgressed ancestry.

Let Λ_*I*_ be the set of 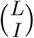 sets of *I* loci among the *L* causal loci, and denote by *λ* a generic element of Λ_*I*_. Let 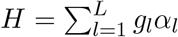 be the haploid genetic value of a gamete. Then, from the law of total variance,

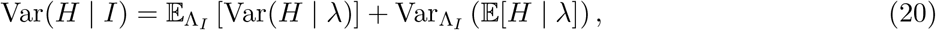

where the subscript Λ_*I*_ indicates that the mean and variance are taken over the (uniform) distribution of the elements *λ* of the set Λ_*I*_. We have

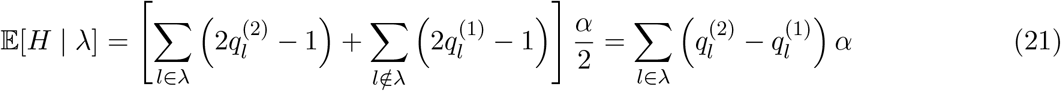

since 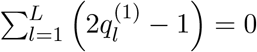, and, ignoring systematically signed linkage disequilibrium among loci within the two populations,

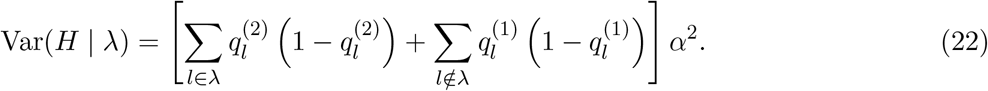

To calculate the first term on the right-hand side of Eq. (20), note that each locus *l* appears in a fraction *p* of the sets *λ* ∈ Λ, i.e., in a number 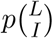 of the sets. Therefore,

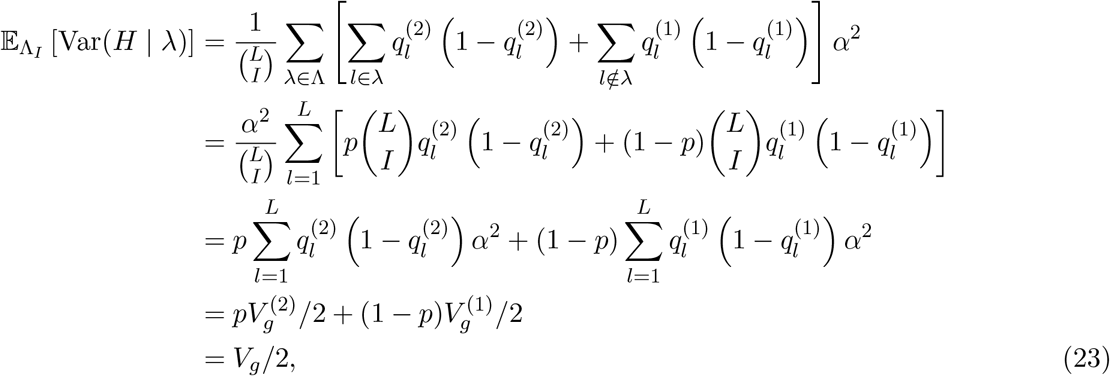

where 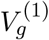 and 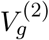 are the genic variances of populations 1 and 2, which we assume to be equal.

To calculate the second term on the right-hand side of Eq. (20), first note that

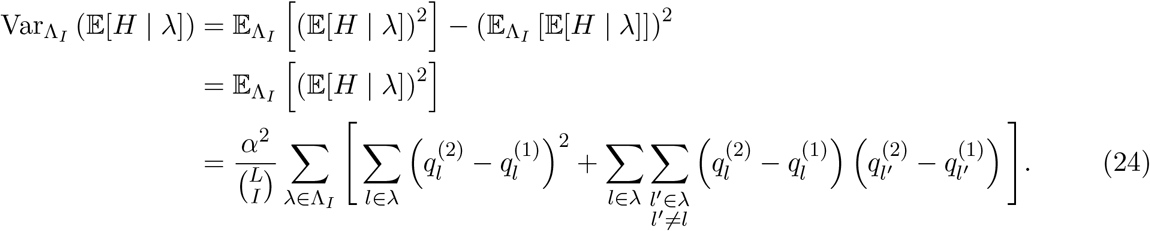

Recall that a given locus *l* appears in 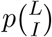 of the sets *λ* ∈ Λ, and note that, for a given pair of distinct loci *l* and *l*^′^, the number of sets *λ* ∈ Λ_*I*_ that contain both *l* and *l*^′^ is 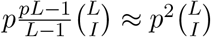. Therefore,

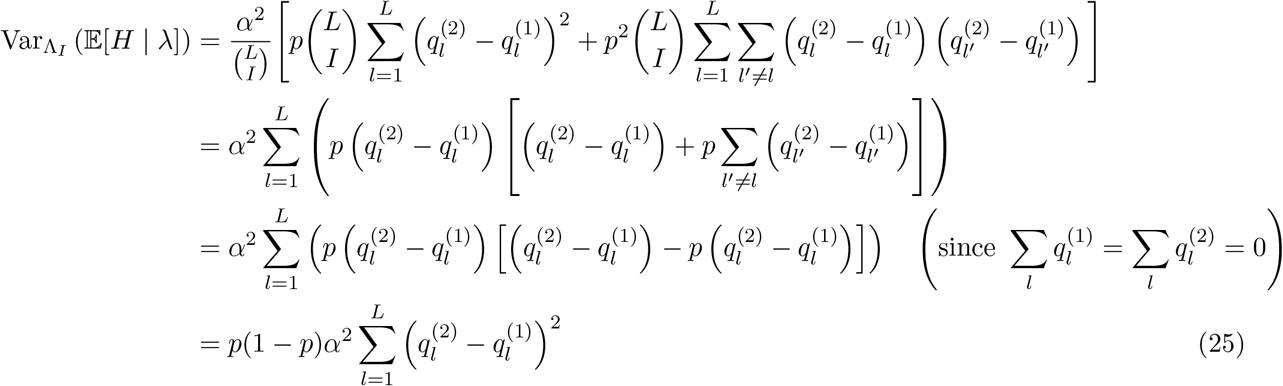

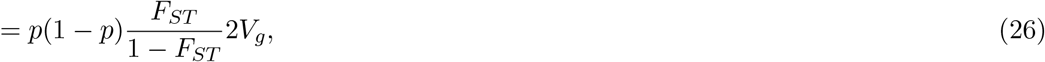

from Eq. (3).

Substituting Eqs. (23) and (26) into Eq. (20), we find that the genetic variance among hybrid gametes with ancestry fraction *p* is

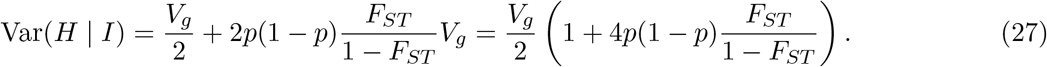

#### Increase in variance for highly diverged populations

The simplification in Eq. (26) assumes that the populations are not highly diverged—indeed, in the limit where *F*_*ST*_ = 1, Eq. (26) is undefined. However, we can use Eq. (25) to calculate the increase in the genetic variance of gametes with ancestry fraction *p* in the case of highly diverged populations that share no genetic variance in the trait (i.e., no shared polymorphisms). The variance of hybrid gametes between these two populations will depend on two sets of loci: (i) loci with fixed differences between the populations and (ii) loci for which one population is fixed and the other is polymorphic.

First, consider the case of fixed differences between the populations. Denote by *L*_*F*_ the set of loci where one population is fixed for a trait-increasing allele and the other is fixed for a (relatively) trait-decreasing allele. At any locus 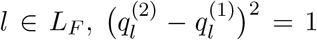, and so the increment to the variance of gametes with ancestry fraction *p* that comes from loci with fixed differences between the populations is, from Eq. (25),

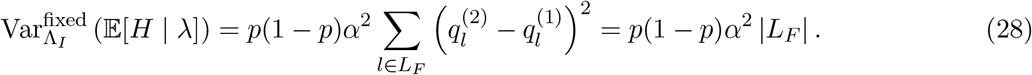

Second, consider the set of loci *L*_*P*_ at which one population is polymorphic and the other is not. Denote by *π*_*l*_ the frequency of the allele at locus *l* that is private to the population harboring the polymorphism at *l*. If the private allele is in population *i* and is trait-increasing, then population *j*≠ *i* is fixed for the trait-decreasing allele, so that 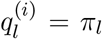 and 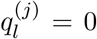. If the private allele is in population *i* and is instead trait-decreasing, then population *j ≠ i* is fixed for the trait-increasing allele, so 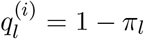 and 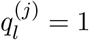. In either case, for the two populations *i* and *j*, 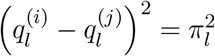. Therefore, from Eq. (25), the increment to the genetic variance among gametes with ancestry fraction *p* that is due to loci for which only one population is polymorphic is

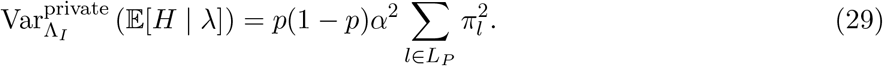

Taking into account both sets of loci—those with fixed differences between the populations and those for which only one population is polymorphic—the genetic variance among gametes with ancestry fraction *p* is

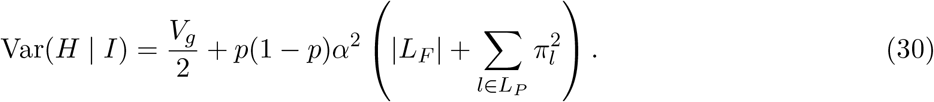

### Long-range LD effects on genetic variance

Denote by 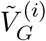 and 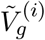 the genetic and genic variance of a gamete sampled from population *i*, with the tilde indicating that these are variances in gametes, not individuals. Under random mating within each population, 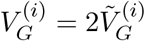 and 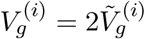. We have

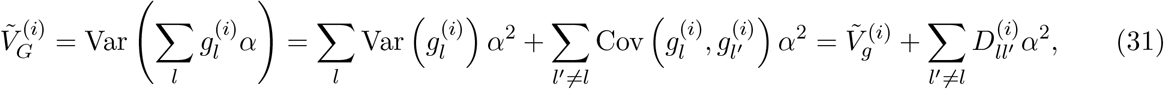

where 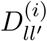 is the coefficient of linkage disequilibrium between the trait-increasing alleles at loci *l* and *l*^′^ within population *i*. We assume that the gametic genetic and genic variances of the two populations are equal to common values 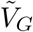 and 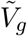, which in turn implies that the values of 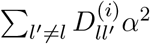 are equal for the two populations, to a common value that we write as 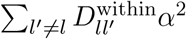.

Consider the set of gametes in which a fraction *p* of the DNA derives from population 2 and a fraction 1 − *p* derives from population 1; under weak selection and free recombination, each such partition is equally likely. In the section above, we show that the genic variance among such gametes is

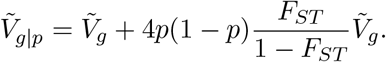

The genetic variance among such gametes is then

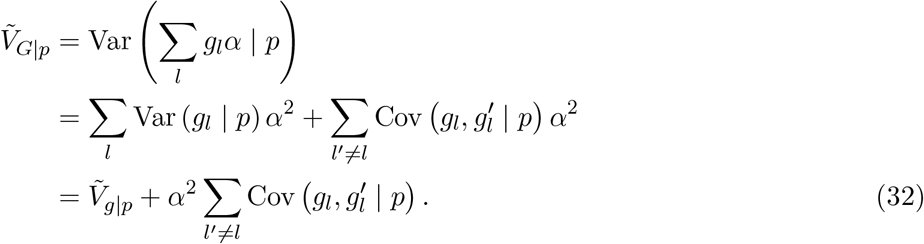

We calculate the covariance term using the law of total covariance:

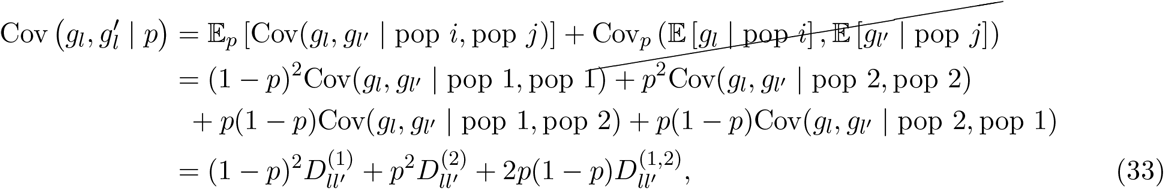

where 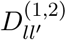 is the covariance in allelic state at *l* and *l*^′^ across the two populations, which we also write 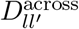. Substituting into Eq. (32),

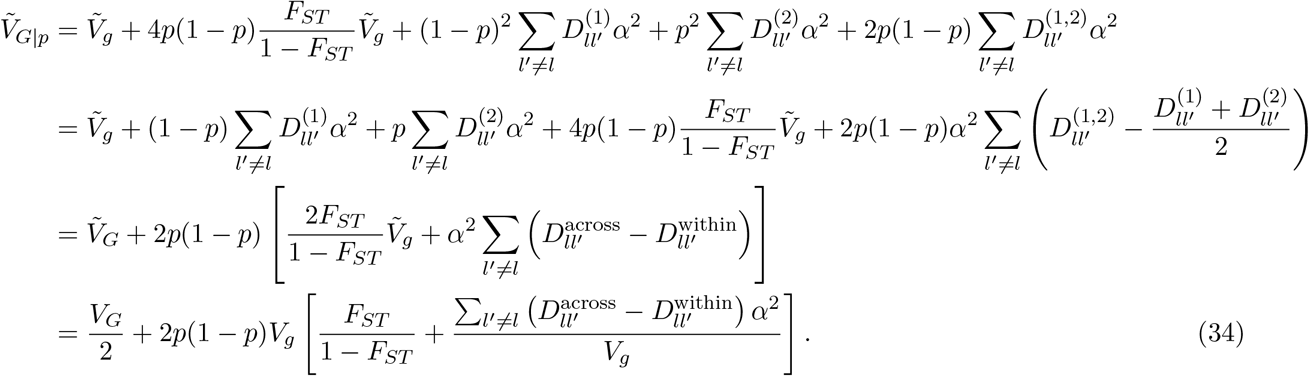

We see that there is a correction to the 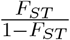 term that depends on the relative contribution of long-range LD to the trait’s genetic variance and on patterns of cross-population long-range LD. Like 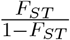, this additional term will go to zero when *F*_*ST*_ goes to zero, because without divergence the two populations would be identical in their long-range LD. In addition, since the relative contribution of long-range LD to trait variance is usually small, this additional term presents only a small correction.

### Expected change in ancestry fraction per generation

We want to calculate the expected change in ancestry fraction per generation, given by the equation

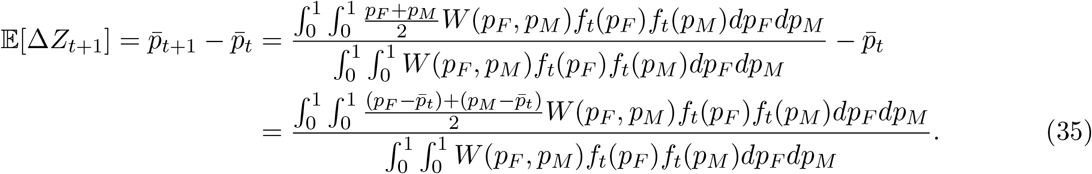

The denominator equals 1 up to a correction of order *V*_*g*_/*V*_*S*_, and we can therefore set it to 1 as we are interested in first order in *V*_*g*_/*V*_*S*_. Therefore, since *p*_*F*_ and *p*_*M*_ are independent and identically distributed with mean 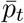

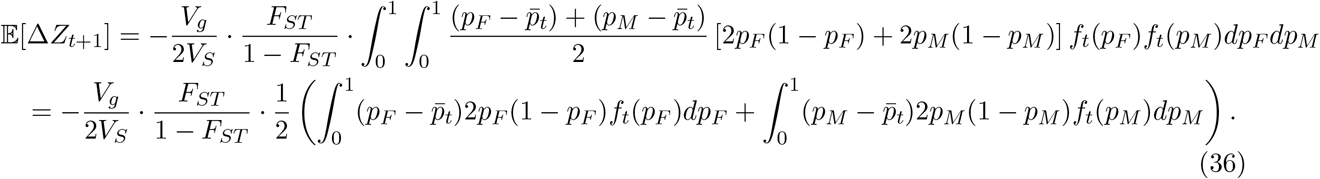

The full Taylor series of *p*_*F*_ (1 − *p*_*F*_) around 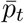 gives the identity

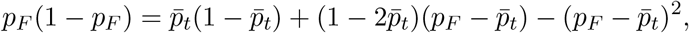

and therefore

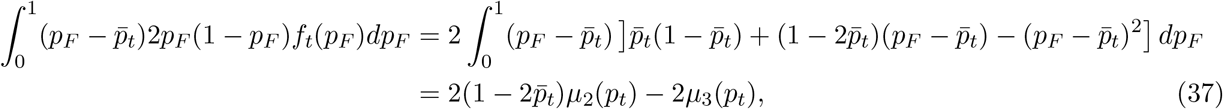

where *µ*_2_(*p*_*t*_) and *µ*_3_(*p*_*t*_) are the second and third central moments of *p*_*F*_ and *p*_*M*_. Similarly,

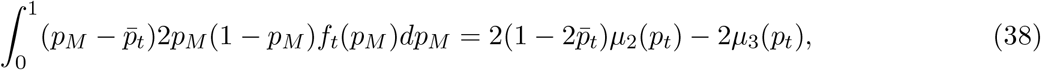

and so

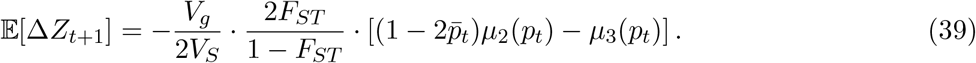

### Simulations

All simulations were carried out in SLiM 4.0 (Haller and Messer 2023). Code is available at https://github.com/cveller.

#### Parameters

We begin by defining a set of loci {1, 2, …, *L*} that are polymorphic in an ancestral population and affect the trait under stabilizing selection. In the simulations displayed in the Main Text, *L* = 1,000. For each locus *l* ∈ {1, 2, …, *L*/2}, we choose an ancestral minor-allele frequency 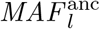. In the simulations displayed in the Main Text, 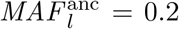 for all loci. At even-numbered loci, the minor allele is trait-increasing, while at odd-numbered loci, the minor allele is trait-decreasing. The effect size at locus *l* is *α*_*l*_: in the simulations displayed in the Main Text, *α*_*l*_ = 1 for all loci. We define 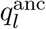 as the ancestral frequency of the trait-increasing allele at locus 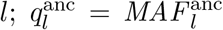 for even *l* and 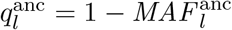 for odd *l*.

Having chosen the ancestral frequencies of trait-increasing alleles 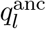, we then choose frequencies for the descendent populations 1 and 2 to ensure a desired value *F* of the *F*_*ST*_ between the two populations, according to the scheme describe below. In the simulations displayed in the Main Text, *F* = 0.2. We opted to sampled allele frequencies for the two populations, rather than having the populations diverge in our simulations independently from the ancestral population until the desired *F*_*ST*_ value was achieved, for the sake of efficiency.

#### Balding–Nichols sampling scheme for allele frequencies

Our aim is to generate, for each locus *l* ∈ {1, 2, …, *L*}, minor-allele frequencies 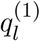 and 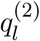 for populations 1 and 2 such that (i) the value of *F*_*ST*_ between the two populations across the set of loci is *F*, in expectation, and (ii) the average trait value in the two populations equals the optimal value of zero (so that there is no immediate directional selection upon admixture, either in favor of one ancestry or the other, or on the average trait value, which might obscure selection against the minor ancestry). Following standard convention, we define the value of *F*_*ST*_ across loci as the average reduction in heterozygosity due to allele-frequency differences between the subpopulations, divided by the average population-wide heterozygosity:

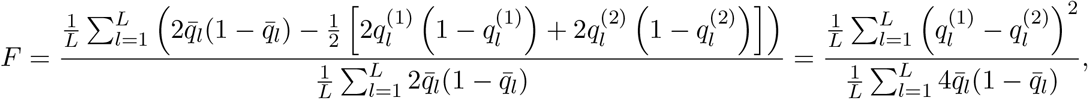

where 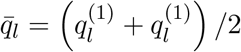. This definition is the one that pertains to our calculations, and is in distinction to the alternative definition of *F*_*ST*_ where we calculate the proportionate reduction in heterozygosity for each locus and then average across loci.

Assuming *L* to be even, for the first *L*/2 loci we draw allele frequencies 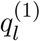 and 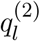 for populations 1 and 2 independently from a beta distribution with shape parameters 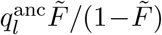 and 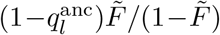, where 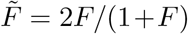. The mean and variance of this distribution are 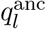 and 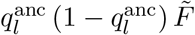, resulting in an asymptotic value of *F*_*ST*_ across the *L*/2 loci of *F*.^1^

Given the frequency 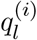 for the trait-increasing allele at locus *l* in the first half of loci {1, 2, …, *L*/2} in population *i*, we then choose the frequencies of the trait-increasing alleles locus 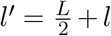 in the second half of loci 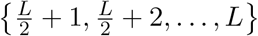 to be 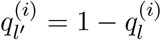. This ensures that, in each population *i*, the mean effect on the trait of locus *l* exactly cancels the mean effect of locus *l*^′^, so that the mean trait value of each population is exactly zero to begin with.

For each set of frequencies sampled in this way, we checked that the realized value of *F*_*ST*_ was within 0.5% of the desired value, and that the genic variances of the two populations, 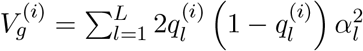, were within 0.5% of each other (so that there is no strong selection in favor of one ancestry or the other to begin with because of lower variance); if either condition failed, we resampled allele frequencies.

#### The strength of selection

We pre-defined a target value of *V*_*S*_/*V*_*g*_ (a value of 8 was chosen for the simulations displayed in the Main Text). In each trial, having sampled allele frequencies as described above, we calculated the average value of 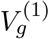 and 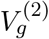, and chose the value of *V*_*S*_ that would govern the strength of selection in that trial by multiplying this average genic variance by our target value of *V*_*S*_/*V*_*g*_.

#### Structure of the genome

Having chosen allele frequencies at the *L* causal loci, we then constructed the genome. We laid down 10,000 loci at which to track ancestry fractions in the eventual admixed population. The *L* = 1,000 causal loci were placed evenly amongst this overall set of loci, such that a causal locus coincided with every 10th locus in the overall set. In each population *i*, the frequency of the trait-increasing allele at the *n*th ordered causal locus was chosen randomly without replacement from the *L* frequencies 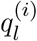: this procedure essentially permutes the frequencies randomly across the *L* causal loci so that, for instance, the sign of the effect of minor allele does not alternate regularly for adjacent causal loci in the genome.

For the case of no linkage (Fig. 3 in the Main Text), the recombination rate was set to 0.5 between ‘adjacent’ loci among the 10,000 total loci. For the case of the human and Drosophila melanogaster linkage maps (Fig. 4 in the Main Text), the 10,000 loci were apportioned among chromosomes according to their physical (Mb) size (as reported in build 38 of the human reference genome^2^ and in release 6 of the *D. melanogaster* reference genome^3^). The loci apportioned to each chromosome were placed evenly along the chromosome’s physical map, and the recombination distances between adjacent loci in each sex were calculated by interpolation of the human male and female linkage maps produced by Kong et al. (2010) and the *D. melanogaster* female linkage map produced by Comeron et al. (2012), using Kosambi’s map function to convert map distances to recombination rates. There is no crossing over in *D. melanogaster* males, and so the recombination rate between adjacent loci on the same chromosome was set to 0. When, in the ordering of the 10,000 overall loci, a pair of ‘adjacent’ loci lay on separate chromosomes, the recombination rate between them was set to 0.5. The resulting 9,999 recombination rates among adjacent loci for human and *D. melanogaster*, for female and male meiosis, were read into SLiM via.`txt` that are available at https://github.com/cveller.

#### Burn-in period

Having defined the structure of the genome, and the frequencies of trait-increasing alleles at the *L* = 1,000 causal loci in each population, we created the two populations, each of size *N* = 10, 000 diploid individuals. We assigned trait-increasing alleles randomly to haploid genomes at each locus in each population according to the frequency of the trait-increasing allele chosen for that population at that locus by the scheme described above, independently across loci, such that, within each population, genotypes would be at Hardy-Weinberg frequencies and there would be no linkage disequilibrium across loci, in expectation.

We then allowed the populations to evolve under stabilizing selection for 20 generations. This burn-in period is long enough to allow the linkage disequilibria that stabilizing selection generates to approach their equilibrium values, and short enough that allele frequencies would not change enough to substantially affect the value of *F*_*ST*_ between the two populations.

#### The admixed population

After 20 generations, we assigned neutral ancestry-marker mutations to all loci in population 2, and mixed the populations (via a single-generation migration event) in the proportions (1 − *θ, θ*) to create a third, admixed population of size *N* = 10,000. A value of *θ* = 0.2 was chosen for the simulations displayed in the Main Text.

We then allowed the admixed population to evolve under stabilizing selection for 500 generations, tracking the frequencies of the ancestry-marker mutations (i.e., the fraction of population-2 ancestry) at each locus.

**Figure S1:**
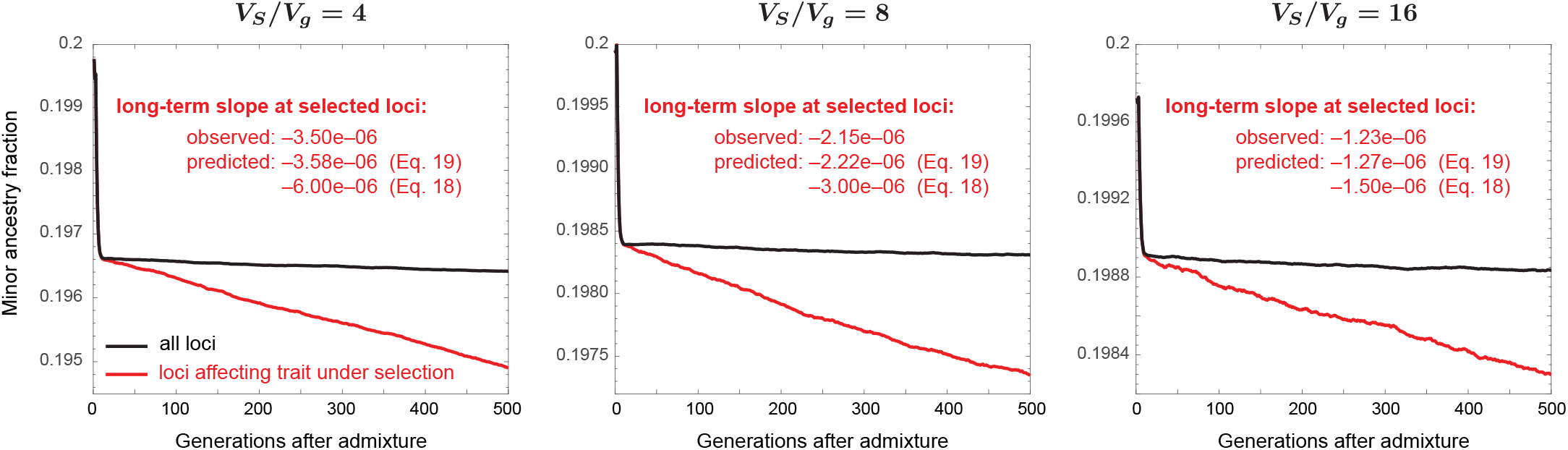
Average minor-parent ancestry over time, with free recombination between all loci, for various strengths of selection *V*_*S*_/*V*_*g*_. The long-term slopes of the trajectories of minor-parent ancestry at trait-affecting loci are calculated from generations 50 to 500, and compared against the predictions of Main Text Eqs. (18) and (19). Other parameter values are as in Fig. 3: *L* = 1,000 causal loci, *θ* = 0.2, *F*_*ST*_ = 0.2. Trajectories are averaged across 1,000 replicate trials. Full details of the simulations can be found in the Methods.

## Footnotes

1 To see this, note that 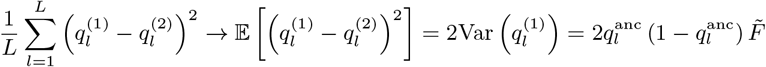 and 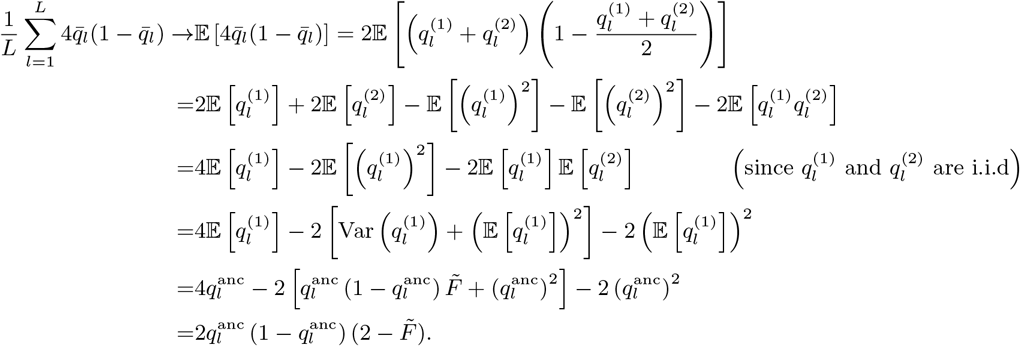 Therefore, 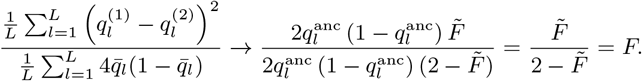

2 https://www.ncbi.nlm.nih.gov/datasets/genome/GCF_000001405.26/

3 https://www.ncbi.nlm.nih.gov/datasets/genome/GCA_029775095.1/

